# Zika Virus Outbreak, Barbados, 2015 – 2016

**DOI:** 10.1101/279182

**Authors:** Sadie J. Ryan, Catherine A. Lippi, Colin J. Carlson, Anna M. Stewart-Ibarra, Mercy J. Borbor-Cordova, Moory M. Romero, Shelly-Ann Cox, Roché Mahon, Adrian Trotman, Leslie Rollock, Marquita Gittens-St. Hilaire, Desmond King, Stephen Daniel

## Abstract

Barbados is a Caribbean island country of approximately 285,000 people, with a thriving tourism industry. In 2015, Zika spread rapidly throughout the Americas, and its proliferation through the Caribbean islands followed suit. Barbados reported its first confirmed autochthonous Zika transmission to the Pan American Health Organization (PAHO) in January 2016, a month before the global public health emergency was declared. Following detection of suspected Zika cases on Barbados in 2015, 926 individuals were described as suspected cases, and 147 lab confirmed cases were reported through December 2016, the end of the most recent epidemiological year. In this short report, we describe the epidemiological characteristics of 926 clinical case records which were originally suspected as cases of Zika, and which were subsequently sent for testing and confirmation; 147 were found positive for Zika, using RT-PCR methods, another 276 tested negative, and the remaining 503 were either pending results or still in the suspected category. Women were represented at about twice the rate of men in case records where sex was reported (71.9%), and confirmed cases (78.2%), and 19 of the confirmed positive cases were children under the age of 10.

---

Zika virus (ZIKV), a Flavivirus transmitted primarily by *Aedes aegypti* and *Ae. albopictus* mosquitoes, was first reported outside of Africa and Asia in 2007. However, it was not until 2015 that Zika rapidly spread from Brazil throughout the Americas. Initially regarded as a mild febrile illness, the emergence of associated health complications such as Zika congenital syndrome (ZCS), including microcephaly and other birth defects, and Zika-associated Guillain-Barré syndrome (GBS), has posed an unprecedented challenge to global health^1,2^. Echoing the rapid spread throughout mainland South America, Zika reached the Caribbean early in the pandemic. Autochthonous transmission in Martinique was first reported in epidemiological week (EW) 51 of 2015, the first case from Puerto Rico was reported in EW 52 of 2015^3^, and many other islands began reporting cases early in 2016^4^. However, case data from several countries has yet to be consolidated and described outside of reports by the Pan American Health Organization (PAHO).

Suspected clinical Zika virus disease cases in Barbados were defined using clinical guidelines provided by PAHO, which include a rash plus one or more of: fever ≥ 38.5°C, conjunctivitis, arthralgia, myalgia, peri-articular edema^6^, but laboratory testing of suspected arboviral cases was also conducted during the Barbados Zika outbreak. Active surveillance of Zika cases (suspected and confirmed) among persons who visited health clinics started as early as May 2015, and the first laboratory confirmed autochthonous case of Zika was reported to PAHO in EW 1 of 2016. However, there were three cases from December, 2015, which were later lab confirmed, of which only one had travel history during the month of infection. Therefore asymptomatic cases may have existed prior to December 2015. Initial Zika case confirmation was conducted using CDC Trioplex RT-PCR assay at for DENV, CHIKV, and ZIKV, at the Caribbean Public Health Agency (CARPHA) laboratory in Trinidad and Tobago, until RT-PCR using the CDC Trioplex assay was established in September 2016 at the *Leptospira* Laboratory, the national reference laboratory of the Ministry of Health of Barbados. Initial testing was biased towards women, particularly pregnant women, reflecting a targeted response. Once testing capability and capacity became local, all samples with suspected arboviral infection were tested for Zika, chikungunya, and dengue viruses. We collected data from records at the Ministry of Health, Barbados, on patients’ age, sex, date of illness onset, occupation, and laboratory diagnostic status (suspected, negative, positive, pending testing). Our reported total suspected case records comprise clinically suspected Zika virus cases prior to September of 2016, and all cases tested as suspected for any of the three arboviral infections, after local testing capacity was established.

The first confirmed Zika case in Barbados, a 42-year-old man, reported onset on December 26, 2015, during EW 51. New cases were subsequently recorded through December 30, 2016 (Fig. 1), with the last cases in 2016 recorded in EW 52. In total, 926 cases with Zika status (suspected, negative, positive, pending testing) were recorded in Barbados in 2015 and 2016, after the first confirmed Zika case, of which 147 (15.9%) were positively confirmed with RT-PCR, and 276 tested negative. The remaining cases were in suspected and pending status at the time of this analysis.

**Figure 1:**
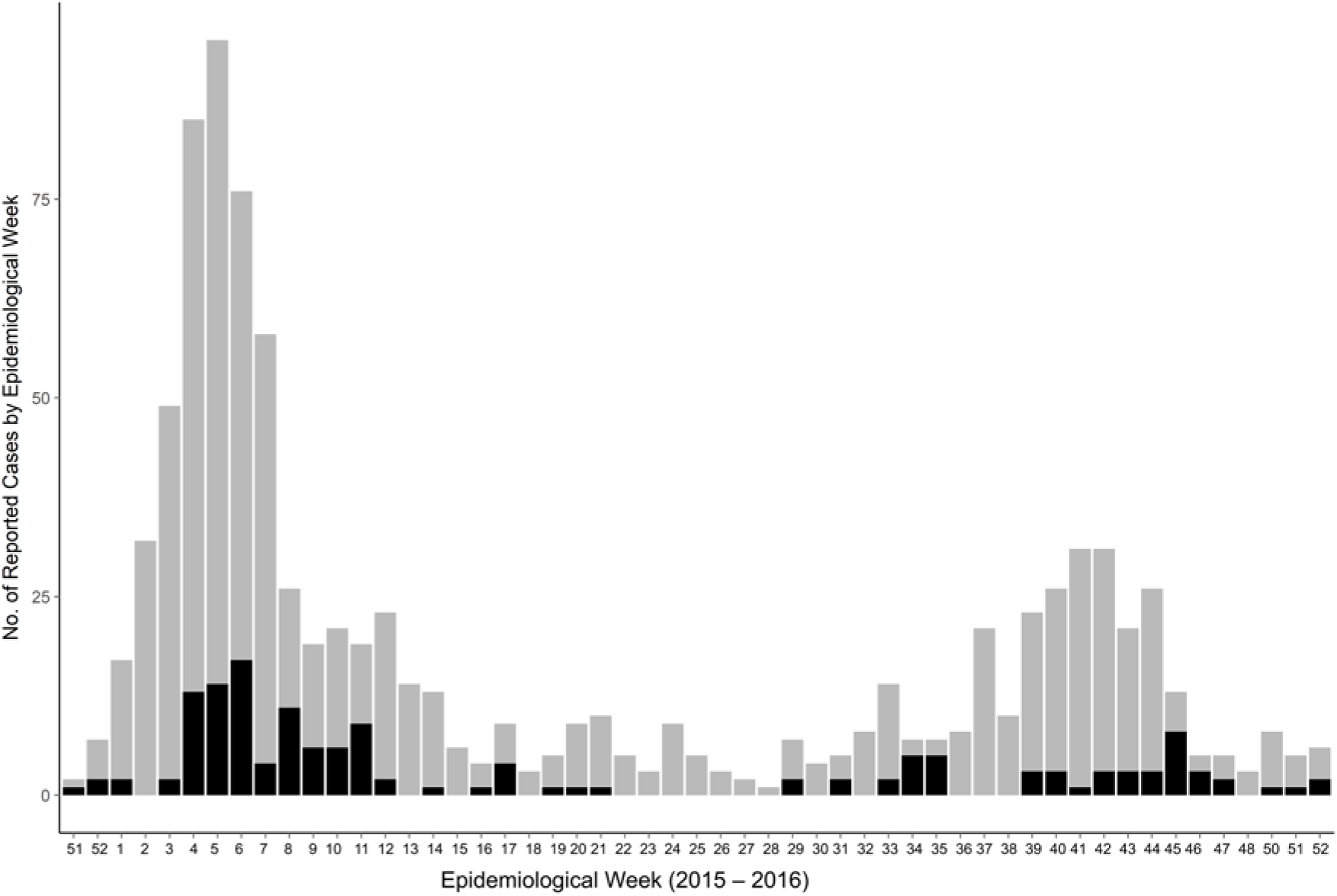
Zika cases (suspected in grey, confirmed in black) reported on Barbados (2015 – 2016).

Sex was reported for 899 of the 926 overall cases in this study (97.1%). Women were disproportionately represented at over twice the frequency of men, with 646 women (71.9%) and 253 men (28.1%). In the confirmed positive cases, this also bore out with 115 women (78.2%) and 32 men (21.8%) in the positively confirmed cases, consistent with a female bias found in other reports on Zika outbreaks^6^. Age was reported for 875 of 926 overall cases (94.5%), with a mean age of 33, and median age of 32; in the positively confirmed cases, mean age was 30, and median of 31, a difference that was not significant. Of the 926 overall cases, 147 (17.0%) were children under the age of ten, and 573 (65.4%) were of childbearing age (15-49). Women of childbearing age represented 77 (52.3%) of the 147 confirmed positive cases, and made up a sizable proportion (48.6%) of the 875 overall cases for which age and sex were both reported. For the 147 positive cases in which age was reported, 19 were children under the age of ten. One of 116 unique occupations were reported for 283 records. We grouped these into five major categories: educational (53), service/hospitality (88), health sector (17), administrative/professional (93), and other (32). The most numerous unique occupational descriptions among these were students (39 plus 2 student nurses), teachers (13), nurses (10), and unemployed (15). The high number of unique occupational descriptions reported, and the low sample of recorded occupations precludes rigorous statistical inference of occupational hazards. The testing status for other arboviral infections for the 926 clinical cases examined for Zika are given in Table 1. It is important to note that dengue testing is conducted at the local Dengue Laboratory, in which a blood sample from suspected dengue cases is sent to the lab from the clinic, and NS1, IgM and IgG, if the sample is from the first 5 days of illness, and if NSI and IgM are negative within the first 3 days, another test is conducted for IgM and IgG after 5 days of illness^7^. In these suspected Zika case records, we cannot distinguish between the trioplex results and blood test results as part of normal surveillance for Dengue. We therefore have far more information about dengue status than for chikungunya. Of the 926 cases in this study, 314 were positive for dengue, and 3 for chikungunya, with an additional 75 suspected for chikungunya. Interestingly, there were 15 positively confirmed Zika-dengue coinfections, but none of the 3 reported confirmed chikingunya cases were coinfections. Other factors of interest when reporting Zika, such as pregnancy status and access to medical care, were not included in the available data for this report.

**Table 1:**
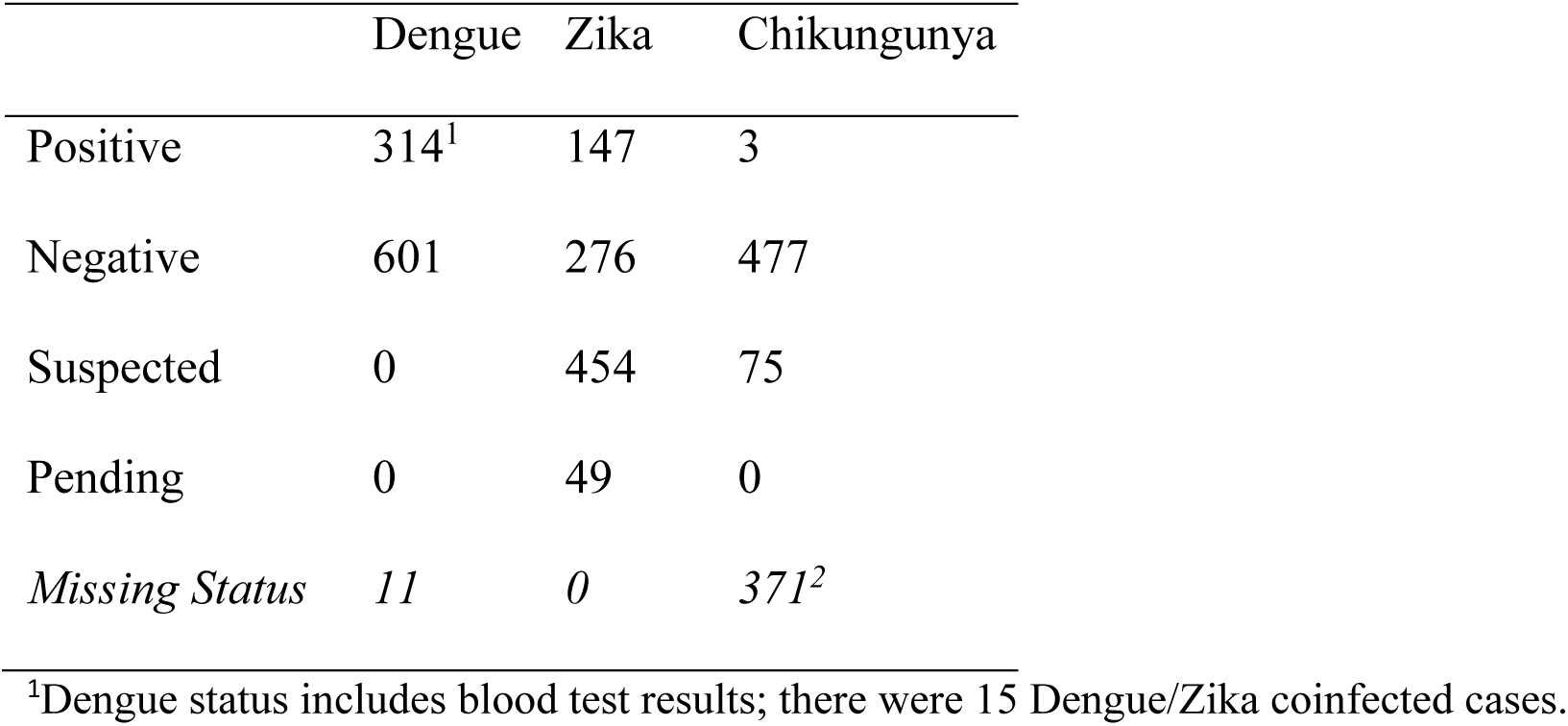
Summary of arboviral infection status (Dengue, Zika, Chikungunya) reported in the 926 total case records.

*Aedes* mosquitoes are established throughout the Caribbean, with active transmission of dengue, chikungunya, and now Zika viruses documented on many islands. In the broader context of emerging arboviruses, the early and rapid onset of the Zika outbreak in Barbados relative to the larger pandemic in the Americas demonstrates that the existence of *Aedes* populations leave even small islands highly susceptible to the spread of novel pathogens. We saw a female bias in cases, particularly toward women of childbearing age, and what appeared to be two waves of cases in 2016 (Fig. 1). The rapid proliferation of Zika infections calls attention to the need to strengthen local capacities for targeted vector control, integrated strategies such as campaigns for cleaning reservoirs, particularly underground cisterns, and health education through formal and informal education programs. In addition this calls for global efforts to support the development of effective vaccines, and a better understanding of the role of sexual transmission and heightened risk to vulnerable populations such as pregnant women.

## Acknowledgements

We thank the Caribbean Institute for Meteorology and Hydrology (CIMH), the people of Barbados, and particularly the Ministry of Health of Barbados. In response to the emerging burden of *Aedes aegypti* transmitted diseases in the Caribbean, the Caribbean Institute for Meteorology and Hydrology (CIMH) has partnered with an international team of researchers to investigate the eco-epidemiology and climate drivers of dengue fever, chikungunya, and Zika fever. This was done through the United States Agency for International Development’s (USAID) Programme for Building Regional Climate Capacity in the Caribbean (BRCCC Programme), executed by the World Meteorological Organization (WMO), and implemented by CIMH. This initiative has brought together the national and regional health and climate sectors (Barbados Ministry of Health, Barbados Meteorological Services, Caribbean Public Health Agency (CARPHA), PAHO and CIMH) to co-develop climate services for the health sector, with a focus on climate-driven spatio-temporal models to predict disease outbreaks.

## Author Contact Information

Sadie J. Ryan and Catherine A. Lippi, Department of Geography and Emerging Pathogens Institute, University of FL, Gainesville, FL. Colin J. Carlson, Department of Environmental Science, Policy, and Management, University of California, Berkeley, Berkeley, CA. Anna M. Stewart-Ibarra and Moory M. Romero, Center for Global Health and Translational Science, State University of New York (SUNY) Upstate Medical University, Syracuse, NY. Mercy J. Borbor-Cordova, Facultad de Ingeniería Marítima, Ciencias Oceánicas y Recursos Naturales (FIMCBOR), Escuela Superior Politécnica del Litoral (ESPOL), Guayaquil, Ecuador. Shelly-Ann Cox, Roché Mahon, and Adrian Trotman, Caribbean Institute for Meteorology and Hydrology (CIMH), Bridgetown, Barbados, and. Leslie Rollock, Desmond King, and Steven Daniel, Ministry of Health, Frank Walcott Building, St. Michael, Barbados, and. Marquita Gittens-St. Hilaire, University of the West Indies at Cave Hill, Faculty of Medical Sciences, Bridgetown, St. Michael, Barbados.

